# Targeted drug release from stable and safe ultrasound-sensitive nanocarriers

**DOI:** 10.1101/2021.12.14.471689

**Authors:** Matthew G. Wilson, Aarav Parikh, Audri Dara, Alexander Beaver, Jan Kubanek

## Abstract

Targeted delivery of medication has the promise of increasing the effectiveness and safety of current systemic drug treatments. Focused ultrasound is emerging as noninvasive and practical energy for targeted drug release. However, it has yet to be determined which nanocarriers and ultrasound parameters can provide both effective and safe release. Perfluorocarbon nanodroplets have the potential to achieve these goals, but current approaches have either been effective or safe, but not both. We found that nanocarriers with highly stable perfluorocarbon cores mediate effective drug release so long as they are activated by ultrasound of sufficiently low frequency. We demonstrate a favorable safety profile of this formulation in a non-human primate. To facilitate translation of this approach into humans, we provide an optimized method for manufacturing the nanocarriers. This study provides a recipe and release parameters for effective and safe drug release from nanoparticle carriers in the body part specified by focused ultrasonic waves.

## 1 INTRODUCTION

Pharmacological treatments are often curbed by intolerable side effects or low effectiveness of drugs (1–7). Consequently, millions of patients remain resistant to treatments and suffer from poor quality of life. There is a critical need for approaches that deliver medication selectively into the desired target in the body at a high concentration while sparing surrounding tissues.

Pioneering work in this domain utilized temperature-sensitive liposomes (8) that can be activated by heat or radiation. However, localized heating at depth is challenging to achieve safely and in a controlled manner. Due to this issue, recent efforts have shifted to using low-intensity ultrasound as a safe and practical form of energy for localized drug release. Ultrasound can be remotely focused into a biological target at depth. Impacting circulating drug carriers, ultrasound triggers targeted release with minimal off-target effects.

Several groups have shown that ultrasound can trigger drug release from nano-sized structures stabilized with biocompatible polymeric shells (9–21), and we have observed effective release in the brain of non-human primates (22).

These nano-sized structures have been commonly filled with perfluorocarbon (PFC) cores. PFCs are highly inert and bestow the resulting nanodroplets with sensitivity to ultrasound. When exposed to ultrasound, the PFC core has been hypothesized to change phase from liquid to gas, greatly expanding in volume, and thus mediating drug release (13, 23–26). Harnessing this understanding, a majority of previous studies used nanodroplets with PFC boiling points below body temperature (9–14, 16–20, 27). However, successful translation of these nanocarriers has been hampered by their instability. By contrast, PFCs with high boiling points are considered to be highly safe and stable and have been well-studied for use in large quantities (28, 29). Nonetheless, such nanodroplets have provided relatively ineffective release (15, 21). In this study, we sought to identify the ultrasound parameters necessary to trigger drug release from high boiling point nanodroplets. We further sought to investigate the biocompatibility and safety of these stable nanodroplets in a nonhuman primate over a repeated dosing regimen.

In particular, this study assessed whether low-frequency ultrasound could enable drug release from stable, high boiling point drug carriers. According to the prevailing hypothesis, ultrasound delivered at the target induces vaporization of the PFC core (13, 23–25). This vaporization may be related to either ultrasound’s mechanical or thermal effects, which vary depending on frequency. High frequencies are more likely to engage thermal mechanisms by depositing more energy at the target. PFCs with high boiling points are unlikely to be influenced strongly by thermal effects and, thus, are expected to release drugs in response to low-frequency ultrasound more effectively. By contrast, high-frequency ultrasound, which has been used predominantly thus far, is likely to favor low boiling point PFCs. In this paper, we demonstrate that the combination of nanodroplet formulation and ultrasound parameters could provide remotely controlled, effective, and safe drug delivery. We also describe the procedure to produce the nanoparticles and release results consistently.

## 2 RESULTS

### 2.1 *In vitro* ultrasound-triggered drug release

We prepared PFC-based, copolymer-stabilized nanodroplets loaded with the neuromodulatory drug propofol (16, 30) and quantified the effectiveness of its release using an established approach in which drug is released from the nanodroplets into an organic solvent (17). Uniquely, we tested how the critical component of the nanodroplet—its core—governs the amount of drug released as a function of ultrasound pressure and frequency. Specifically, we tested the release effectiveness of three different PFC cores—perfluoropentane (PFP), decafluoropentane (DFP), and perfluorooctylbromide (PFOB). These PFCs have boiling points of 29°C, 55°C, and 142°C, respectively. Nanodroplets with these cores had comparable sizes: mean±SD of 543.3±23.7, 550.8±91.7, and 473.0±28.4 nm for PFP, DFP, and PFOB, respectively.

We used an ultrasonic transducer capable of operating at low (300 kHz) and high (900 kHz) frequency focused on vials containing the nanodroplet solutions (31). We found that 300 kHz ultrasound triggered more drug release from the nanodroplets than 900 kHz (Fig. 1, left). The difference in the percentage of drug released at the two frequencies (31.7% and 20.3%, respectively) was significant when averaged across all cores and ultrasound pressures (*t*(130) = 3.3, *p* = 0.0013, two-sample two-tailed t-test). In line with our hypothesis, we found (Fig. 1, right) that at the higher 900 kHz frequency, the release effectiveness strongly depends on the boiling point of the PFC core—the lower the boiling point of the PFC, the higher the release effectiveness. An omnibus ANOVA model (Table S1) that incorporated all factors tested (core, ultrasound frequency, ultrasound pressure) as well as all possible interactions, detected a significant core × frequency interaction (*F* (2, 90) = 8.05, *p* = 0.00061).

**Figure 1.**
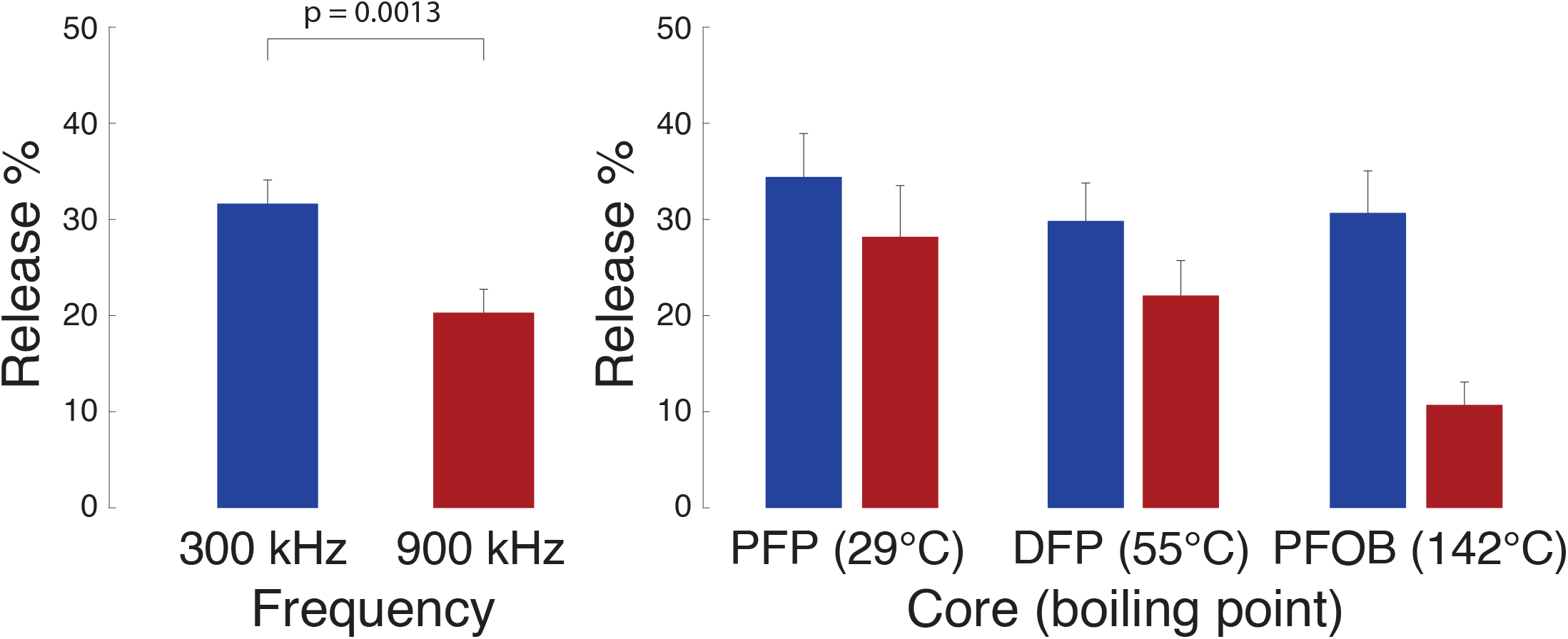
Release from nanodroplets with distinct cores under two ultrasound frequency modes. Mean±s.e.m. propofol release relative to the amount encapsulated (see Materials and Methods for details) for the two ultrasound frequencies (left, combined across all cores) and the three cores (right) tested. The ultrasound was delivered in 100 ms pulses repeated 60 times over the period of 1 minute. The p-value denotes the significance of a two-sample two-sided t-test. A complete statistical analysis of the effects is provided in Table S1.

The scaling of the release effectiveness by the PFC boiling point (red bars in Fig. 1, right) suggests the engagement of a thermal mechanism at the higher frequency, in agreement with previous propositions (13, 18, 24, 32). If this is the case, the release should also scale with the average ultrasound intensity *I* delivered into the nanodroplets. For longitudinal waves, it holds 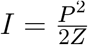 (33), where *P* is the ultrasound pressure amplitude, and *Z* is the acoustic impedance of the medium. Thus, a thermal effect scales with pressure squared. We indeed found that drug release at 900 kHz showed a quadratic dependence on pressure (Fig. 2, right). Specifically, quadratic fits to the data (solid curves) explained 96.5, 93.5, and 94.8% of the variance in the PFP, DFP, and PFOB data points. In comparison, linear fits only explained 80.3, 80.5, and 69.3% of the variance, respectively. The difference in the mean variance explained by the quadratic and linear fits (94.9% versus 76.7%) was significant (*t*(2) = 4.83, *p* = 0.040, two-sided t-test). In contrast, the lower 300 kHz frequency showed a linear dependence of the release on pressure (Fig. 2, left). A quadratic fit did not significantly (*p >* 0.16) differ from the linear fit in terms of variance explained (89.5% versus 89.3%). The linear dependence on pressure at 300 kHz is consistent with a mechanical rather than a thermal effect (see Discussion for details).

**Figure 2.**
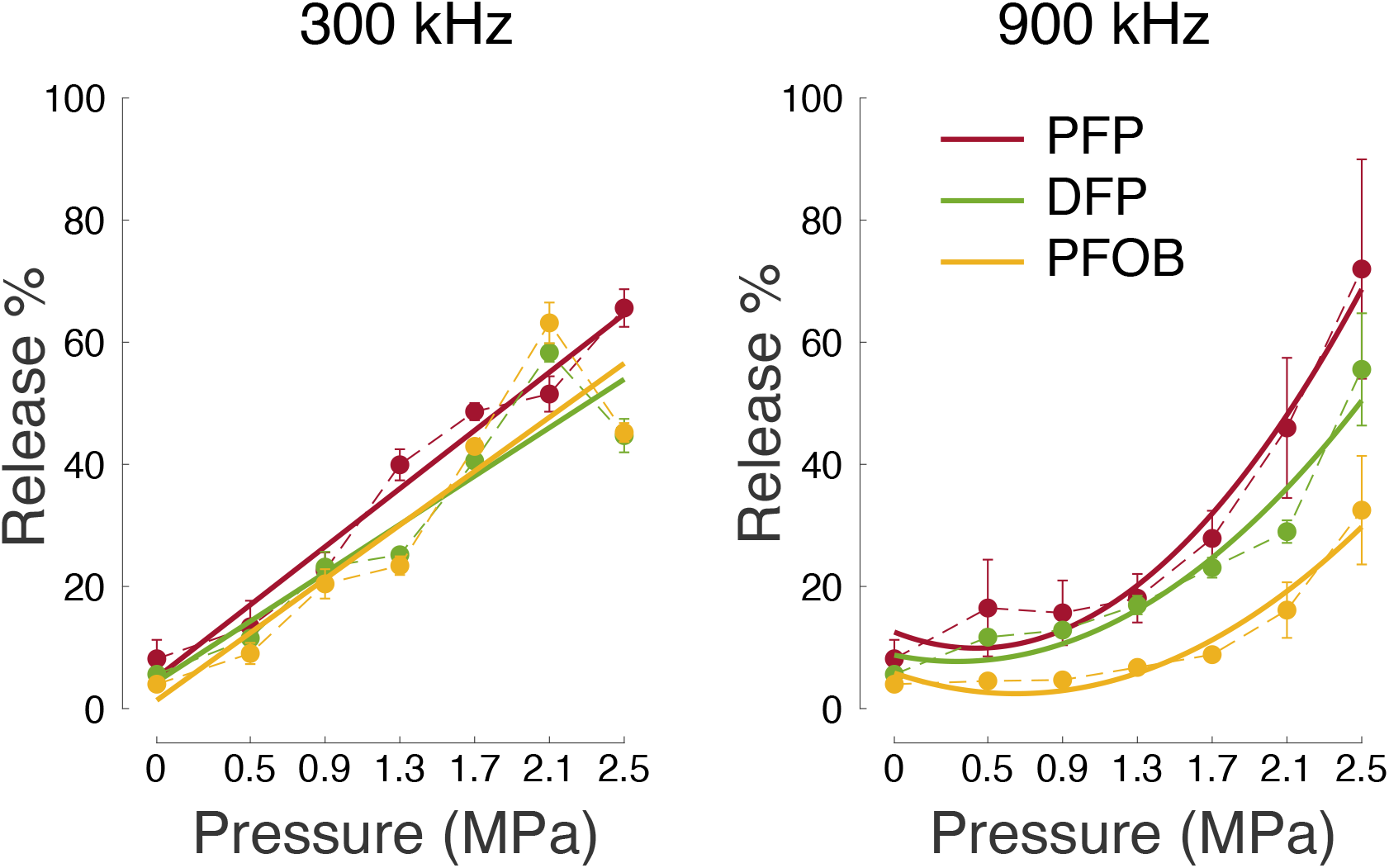
Release across all tested factors. Mean±s.e.m. percentage of the released propofol for the two ultrasound frequencies (left versus right), ultrasound pressure amplitude (abscissa), and the three cores tested (color; see inset). The plots used *n* = 3 samples for each core and ultrasound parameter except for 0 MPa, which used *n* = 4. The thick lines represent quadratic fits to the data. For the 300 kHz data, linear fits were comparably explanatory as the quadratic fits (see text).

To further validate effective release, we have complemented these *in-vitro* data with *in-vivo* release data in the brain of non-human primates, a topic of a dedicated study (22).

We summarize the effects of the three factors tested (core, ultrasound frequency, and ultrasound pressure) as well as all possible interactions in an omnibus ANOVA model (Table S1). This analysis confirms that both the core and ultrasound parameters (frequency and pressure) are significant factors for effective release. In addition, the interaction of the factors indicates that the selection of a specific core must be performed in conjunction with the appropriate ultrasound frequency.

### 2.2 Stability of PFC nanodroplets

From a safety perspective, cores with higher boiling points can be expected to be more stable, thus minimizing the risk of spontaneous drug release and embolism when injected into the bloodstream. Indeed, we found that the low-boiling point, PFP-based nanodroplets more than doubled in size over 24 hours at room temperature (Fig. 3. *t*(5) = 11.9, *p <* 0.001, paired two-tailed t-test). In contrast, the size of PFOB-based nanodroplets remained steady, increasing only by 23 nm over the 24-hour period (*t*(5) = 6.3, *p* = 0.0015). A repeated measures ANOVA detected a highly significant interaction of PFC core and time (*F* (5, 50) = 43.54, *p <* 0.001), confirming that the PFP-based nanodroplets expanded at a significantly higher rate than PFOB-based nanodroplets.

**Figure 3.**
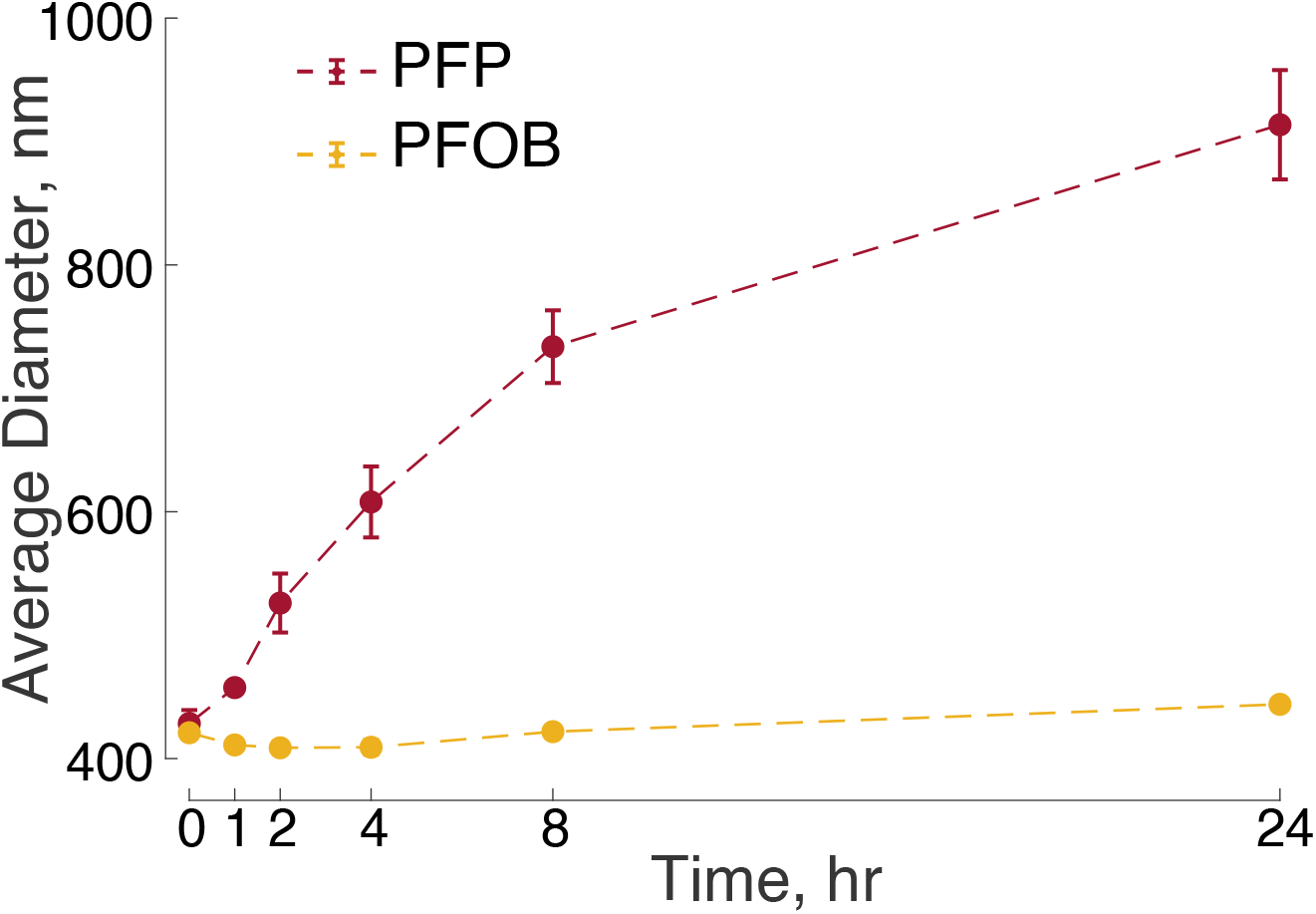
Stability of PFOB and PFP-based nanodroplets over time. Mean±s.e.m. diameter measured at the times as indicated on the abscissa. The times were measured relative to the time of the completion of the nanodroplet production. The data include *n* = 6 samples of PFOB nanodroplets and *n* = 6 samples of PFP nanodroplets. The error bars for PFOB are smaller than the symbols.

### 2.3 Safety

We have evaluated the safety of the PFOB-based nanodroplets in a non-human primate. To evaluate the safety of repeated dosing of PFOB nanodroplets, we completed a series of blood draws to monitor clinical chemistry and hematology parameters. A total of six doses of nanodroplets were administered: an initial ramp-up at 0.25, 0.5, and 1.0 mg/kg of propofol followed by three additional doses at 0.5 mg/kg, which we have previously found to elicit robust effects on choice behavior of non-human primates (22). This approach provides detailed insights into the potential toxicity of the nanocarriers to the liver (ALP, AST, ALT, total protein, albumin, and bilirubin), kidneys (creatinine, BUN), or spleen (RBC, WBC, and platelet counts) where these nanodroplets are expected to accumulate (14, 22). We also assessed the level of immune system activation using the WBC count. These key indicators along with the normal range for rhesus macaques are shown in Fig. 4. All other blood chemistry and hematology markers are shown in Fig. S3. To evaluate both short-term and long-term effects, we quantified changes in these parameters 1 and 7 days after each nanodroplet dose at or above 0.5 mg/kg (Fig. S4). Across these comparisons, only blood glucose showed a significant difference (*t*(4) = *−*6.03, *p* = 0.004; one-sample t-test), decreasing on average by 21 mg/dL 1 day following the injection. The results associated with the individual t-test as well as 95% confidence intervals are shown in Table S3.

**Figure 4.**
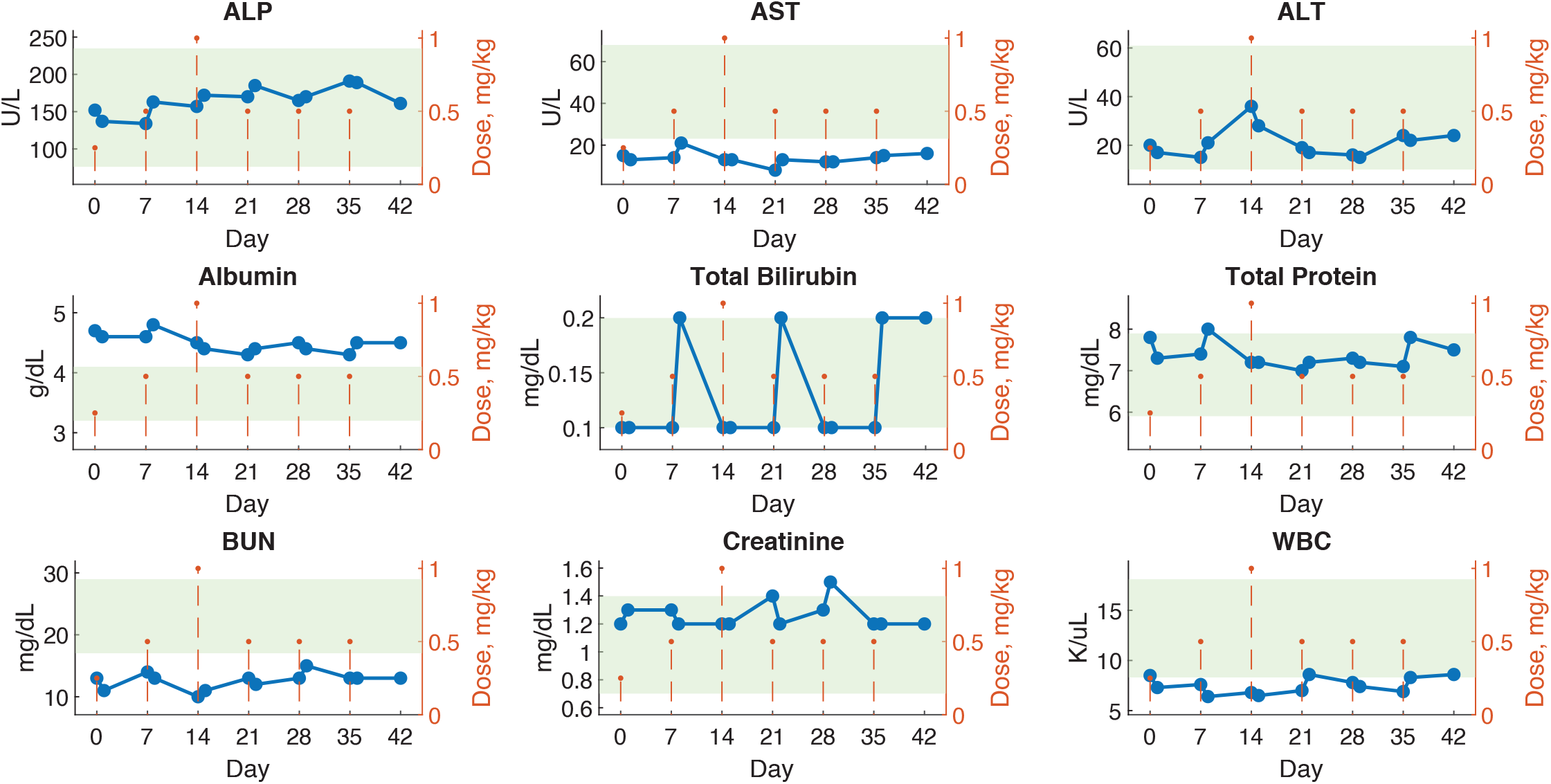
Blood metrics of safety show minimal changes following 6 weeks of continuous nanodroplet administration to a non-human primate. Blood chemistry and hematology values for a macaque monkey plotted over 42 days with 6 nanodroplet doses (orange dotted lines). Blood samples were collected 1 and 7 days after each dose (blue points). Green shaded areas represent normal values established by the Association of Primate Veterinarians. The displayed parameters are metrics of liver, kidney, and immune response function. Fig. S3 shows all other parameters.

### 2.4 A versatile manufacturing process for ultrasound-responsive nanocarriers

To demonstrate the versatility of the PFOB nanodroplets, we loaded them also with ketamine (an anesthetic with longer effect duration) and mycophenolate motefil (an immunosuppressant). We observed a similar sensitivity to ultrasound as the propofol-loaded nanodroplets (Fig. 5). Indeed, an ANOVA model incorporating ultrasound pressure and drug type indicated no significant effect of drug type (*F* (1, 46) = 0.88, *p* = 0.4). Mycophenolate motefil-loaded nanodroplets did release more drug without ultrasound (*t*(5) = 6.5, *p* = 0.0013, two-sample two-tailed t-test). This may be attributed to the lower hydrophobicity compared to propofol and ketamine, leading to less effective encapsulation inside the nanodroplets.

**Figure 5.**
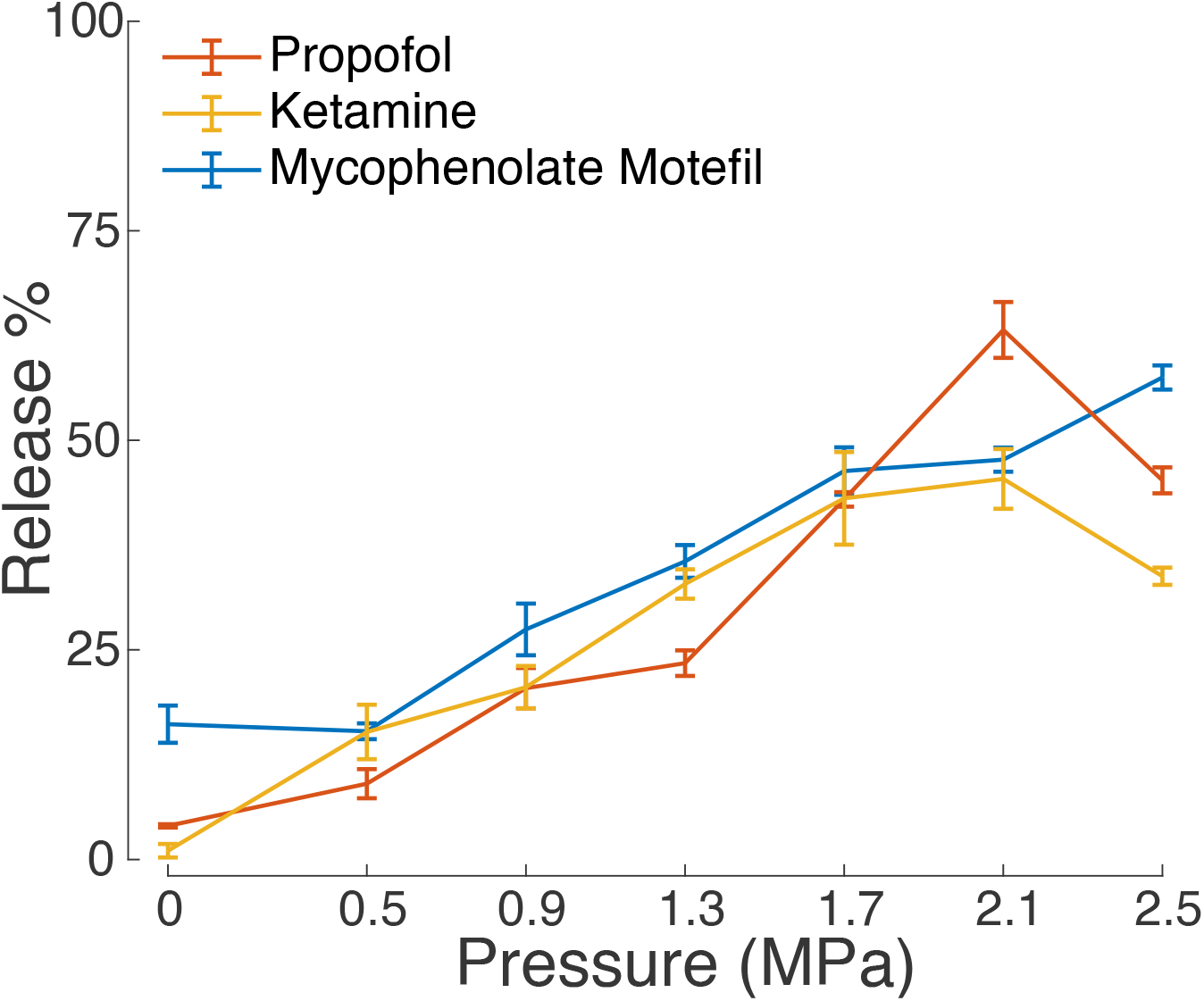
Hydrophobic drugs encapsulated and released by PFOB nanodroplets. Mean±s.e.m. percent drug released under ultrasound stimulation under the same experimental conditions as in Fig. 2.

To provide the reader with a recipe to reliably manufacture the nanoparticles, we systematically studied key variables in the manufacturing process. This included the ratio of nanodroplet components (drug and PFOB to polymer), sonication bath parameters (temperature and duration), and centrifuge parameters (speed and duration). The results are summarized in Fig. 6, which shows how each variable affects drug encapsulation and release. Table S2 summarizes the statistical analysis of associated with this evaluation. The table indicating which manufacturing parameters significantly affect the nanodroplet size, drug encapsulation, and release.

**Figure 6.**
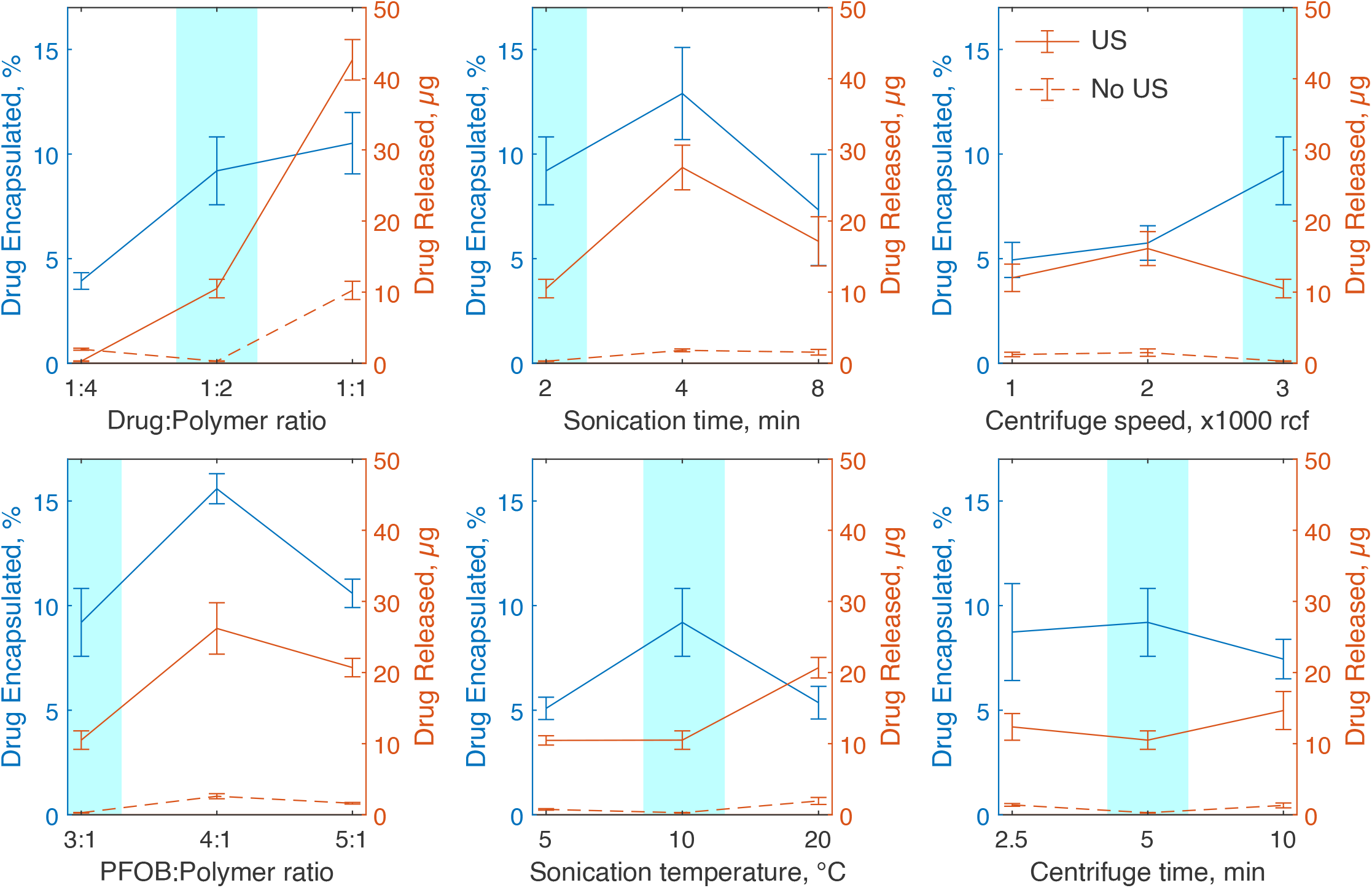
The effects of nanoparticle manufacturing parameters on drug encapsulation and release. Each plot shows the mean±s.e.m. propofol encapsulated as a percentage of propofol used in blue and total drug released with (solid line) and without ultrasound (dotted line). In these tests, 300 kHz ultrasound was applied for 60 seconds in 10 ms bursts at a rate of 10 Hz and peak negative pressure of 1.5 MPa. Each parameter test involved 5 independent nanoparticle samples. Baseline parameters highlighted in blue were held constant while other parameters were varied.

These data show that the ratios of polymer shell to drug and PFOB core strongly influence the resulting nanodroplet composition, and thus must be selected careful to meet specific application criteria. For example, increasing the ratio of drug to polymer shell significantly increases the rate of encapsulation and drug release, but also increases the rate of spontaneous drug release and the average particle size (a change from 780±80 nm to 3300±600 nm in size). A full summary of size results can be found in Supplementary Fig. S1. We found no significant linear correlation between particle size and encapsulation (*p* = 0.61) or particle size and drug release with (*p* = 0.47) or without ultrasound (*p* = 0.89) (Supplementary Fig. S2).

## 3 DISCUSSION

This study investigated the effectiveness and safety of drug release from biocompatible nanodroplets triggered by focused ultrasound. Uniquely, we evaluated the release efficacy from nanodroplets filled with PFCs with distinct boiling points, including boiling points above the body temperature, and investigated the relevant ultrasound parameters that activate them. We found that low frequency ultrasound (300 kHz in particular) was more effective at mediating release in general and that low frequencies are paramount for effective release from PFCs with a high boiling point.

The core has been hypothesized to be a critical factor in governing the effectiveness of ultrasound-based drug release (11, 18, 21). Indeed, our 900 kHz data show (Fig. 1, right, red) that at the range of commonly applied ultrasound frequencies (about 1 MHz and greater), the core is a key determinant of release effectiveness. We found that the higher the boiling point of the encapsulated PFC, the lower the release effectiveness. This finding is in accord with previous results (13, 18, 24, 32). Critically, we found that lowering the frequency to 300 kHz increased the release effectiveness at a general, core-independent level (Fig. 1). This effect can also be found in a study that used PFP (17). The overall effect of decreasing frequency may be enhanced by the increase in focal volume, but the differences between cores at each frequency remain informative. The application of low-frequency ultrasound thus opens the path to using high boiling point cores (e.g., PFOB) as release actuators. A frequency of 300 kHz provides spatial resolution on the order of several millimeters, which approximates the focal volume of the drug released in tissue by ultrasound (16, 19). While the focal size is larger than ultrasound of higher frequencies, the focal size remains applicable for confined targets, including in the brain (34, 35).

This work uniquely evaluated the safety of ultrasound responsive PFC nanodroplets over 6 weeks of repeated dosing in a nonhuman primate. This type of dosing regime may be necessary for long-term treatments. The minimal impact on measures of toxicity in a nonhuman primate over this time period are a key indicator of potential for clinical translation. Indeed, only blood glucose was found to show a statistically significant change at 1 day after the nanodroplet administration. The level stabilized at day 7. This change in blood glucose may be attributed to diet—the monkey received sweet juice as a reward on the days of the nanodroplet administration.

The majority of studies that applied ultrasound to PFC-based nanodroplets used PFCs with boiling point below the body temperature (9–14, 16–20, 27). Although the Laplace pressure of sub-micron particles can effectively increase the boiling point when encapsulated (36), PFCs with low boiling points may suffer from instability issues, which has raised safety concerns. Indeed, our data show that PFP-based nanodroplets not exposed to ultrasound spontaneously increase in size over a short time window, whereas PFOB nanodroplets remain stable (Fig. 3). These data suggest that PFP nanodroplets, if at all, should be used immediately following their production. It is unclear if PFP nanodroplets would vaporize more quickly than they are cleared from the bloodstream, however; more work is required on this matter. Further, low boiling point PFCs can form persistent microbubbles after vaporization by ultrasound (10, 13, 20). Our finding that PFOB can be used for effective release from nanodroplets at low ultrasound frequencies (Fig. 1, Fig. 2) should mitigate the boiling-point associated concerns. Indeed, PFOB-based products including LiquiVent—an oxygen-carrying liquid drug, and Oxygent (both Alliance Pharmaceutical Corporation, San Diego, CA, USA)—a blood substitution agent (28, 29, 37) have been thoroughly studied for use in liter-quantities.

The *in vitro* release data presented here also contribute to the understanding of the release mechanism. Thus far, the predominant hypothesis of action has been the vaporization of the PFC droplet upon the impact of focused ultrasound (11, 18, 21, 23–25). In this mechanism, the thermal and mechanical aspects of propagating ultrasound exceed the vaporization threshold governed by the PFC boiling point and the Laplace pressure. The PFC core increases in size (up to a 5-fold increase in diameter (36)), which contributes, in addition to any superimposed effects of ultrasound, to drug release. Indeed, our data at the common range of ultrasound frequencies, 900 kHz, provide two lines of evidence in support of this hypothesis. First, we found that the release increases with decreasing boiling point (Fig. 1, right), as expected from the vaporization hypothesis. Second, thermal energy delivered by ultrasound is known to be proportional to pressure squared (33). We indeed found a quadratic dependence of the release on pressure (Fig. 2, right).

Our and a previous study (17) suggest that ultrasound of frequencies lower than those in the common diagnostic range may mediate more effective drug release. Lower frequencies are known to accentuate mechanical effects, which can take two forms. First, ultrasound can induce cavitation, the formation of gaseous nuclei from dissolved gasses under the low-pressure epochs of the ultrasound pressure wave (38). Cavitation can provide useful mechanical forces until it exceeds a threshold at which the formed gaseous nuclei collapse and cause damage (38). From the FDA’s 510(k) Track 3 standpoint (39), cavitation is unlikely to occur for mechanical index—defined as ultrasound pressure divided by the square root of frequency— values below 1.9. In our hands, a 300 kHz, 1.0 MPa pressure at target yields a mechanical index of 1.83. Despite being below the value of 1.9, this pressure level already drives significant release (Fig. 2, left). Thus, if cavitation is involved, it likely constitutes one of multiple mechanisms. Another candidate mechanism for drug release is particle displacement. The maximal displacement of a particle in the ultrasound path, *ξ*_*m*_, is linearly proportional to the ultrasound pressure amplitude 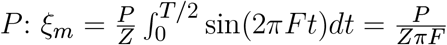, where *F* is the ultrasound frequency and *Z* the acoustic impedance. Indeed, our 300 kHz data show a linear dependence of the release on pressure (Fig. 2, left). This mechanism is supported by high-speed imaging, which did not detect a persistent phase change despite effective release in rodents (17). Together, our findings indicate that both thermal and mechanical effects can be at play, depending on the applied ultrasound frequency. Higher frequencies accentuate thermal effects (PFC vaporization), whereas lower frequencies accentuate mechanical effects (cavitation or cycle-by-cycle particle displacement).

Interestingly, at 300 kHz, we observed a marginal decrease in drug release from DFP and PFOB nanodroplets at the highest ultrasound pressure tested (Fig. 2, left). This effect has been also observed in another study (19). These high pressures may lead to cavitation—the formation of bubbles—which may consequently block the propagation of ultrasound into the target (19). PFP did not exhibit this effect, possibly due to the lower boiling point—complete vaporization and subsequent collapse of PFP bubbles may prevent shielding effects with the repeated pulsing scheme used here.

The study has three limitations. First, we tested the effects of ultrasound frequency and pressure, but not the ultrasound waveform. For instance, we have not tested the effects of the duty cycle and only used a fixed value of 10%. Increases in the duty cycle are known to lead to stronger, cumulative effects (14, 17). Second, we quantified the release of nanodroplets using a single copolymer shell (PEG:PDLLA). This limitation is mitigated by a previous study that found little variability in the release due to specific forms of the polymeric shell (17). And third, the *in vitro* drug release assay does not fully replicate the conditions *in vivo*, which we have evaluated in a dedicated study (22). Blood contains many proteins and cells which may interfere with the function of the nanodroplets or diffusion of drug after activation, and uptake into brain tissue may differ from uptake into hexane. Thus, while the absolute values for drug release may not translate, we here emphasized the relative comparisons between different nanodroplet compositions and ultrasound parameters.

## 4 CONCLUSIONS

In summary, we found that low-frequency ultrasound can effectively release drugs from stable PFC-based nanocarriers. This study informs the use of specific PFC-based nanodroplets and ultrasound parameters for effective, safe, and targeted drug release in humans while providing evidence of biocompatibility in nonhuman primates over an extended period of time. To facilitate the translation into applications in humans, we report the parameters critical for robust assembly and production of the nanoparticles.

## 5 MATERIALS AND METHODS

### 5.1 Materials

Methoxy poly(ethylene glycol)-b-poly(D,L-lactide) (PEG-PDLLA) co-polymers with 2,000 : 2,200 g/mol molecular weights, respectively, were obtained from PolyScitech (USA). 2H,3H-decafluoropentane and perfluorooctyl bromide were obtained from Tokyo Chemical Industry Co. (Japan). Perfluoro-n-pentane was obtained from Strem Chemicals (USA). Propofol was obtained from Sigma Aldrich (Millipore Sigma, Canada). Infrared dye IR800RS NHS Ester was obtained from LI-COR Biosciences (USA). HPLC-grade tetrahydrofuran (THF) and methanol were obtained from Fisher Scientific (USA). Phosphate buffer solution (PBS) was obtained from Gibco (Thermo Fisher Scientific, USA) and Cytiva (USA). Mycophenolate Motefil was obtained from Akorn, Inc. (USA). Ketamine was obtained from VetOne (USA).

### 5.2 Nanodroplet production

The process of manufacturing the drug-encapsulating, ultrasound-responsive PFC particles is illustrated at a conceptual level in (Fig. 7) and described in detail in previous studies (9, 17). The process converts small (*<* 30nm) micelles into much larger (*>* 300nm) PFC-filled nanodroplets. First, the PEG-PDLLA polymer constituting the basis of the nanodroplet shell is dissolved in THF at a rate of 1mL THF : 16 mg polymer. For the biodistribution and blood clearance studies, infrared dye is added at a ratio of 1:32 (dye:polymer) for the rats and marmoset and 1:110 or 1:89 for the macaques 1 and 2, respectively. THF is then evaporated under vacuum until a gel-like layer remains. PBS is added at a rate of 1 mL PBS : 8 mg polymer and placed on a shaker table at 120 rpm to dissolve for 15 minutes. The addition of PBS orients the hydrophilic copolymer, PEG, toward the water and the hydrophobic, PDLLA, copolymer away from the water, and as a consequence, micelles are formed. Next, the PFC core and propofol are added and emulsified. A ratio of 1 mg propofol : 2 mg polymer was used in all cases. The nanodroplets’ diameter can be controlled by the ratio of PFC to polymer, as reported previously (11). For PFOB and DFP nanodroplets, a ratio of 4.5 *μ*L PFC : 1 mg polymer was used. The ratio for PFP was scaled up to 6.25 *μ*L : 1 mg to account for PFC lost to vaporization before being emulsified. A 20 kHz, 500W sonicator with a cup horn attachment (VCX500, Sonics) was used to perturb the thermodynamic equilibrium of the micellar system, which leads to the incorporation of PFOB into the micelles and the formation of stable nanodroplets or nanodroplets (40). The PFC and drug are added to 15 mL centrifuge tubes and gently shaken to combine before adding 8 mL of the micelle solution. The samples are then sonicated in a cold bath at 20% power in 30-second intervals until the solution is cloudy and drug and PFC are fully emulsified (1-3 minutes in total). A custom temperature-controlled cooling system maintained the bath temperature during sonication at 2°C for PFP and 10°C for DFP and PFOB. PFP must be kept colder to minimize vaporization before emulsification, while DFP and PFOB require higher temperatures to emulsify successfully without vaporizing. We found this controlled temperature approach to maximize the consistency of the nanodroplet sizes, drug encapsulation, and release properties. The resulting solution contains the desired nanodroplets in addition to remaining micelles, dissolved polymer, and free propofol. Nanodroplets are isolated using three cycles of centrifugation at 3,000 relative centrifugal force (RCF) at 4°C. After each cycle, the supernatant is discarded and the pellet dissolved in 5 mL fresh PBS. If the resulting solution contains larger particles than needed, these were removed by a slower centrifuge cycle for 1 minute at 800 RCF, this time keeping the supernatant. Larger particles contained in the pellet are discarded.

**Figure 7.**
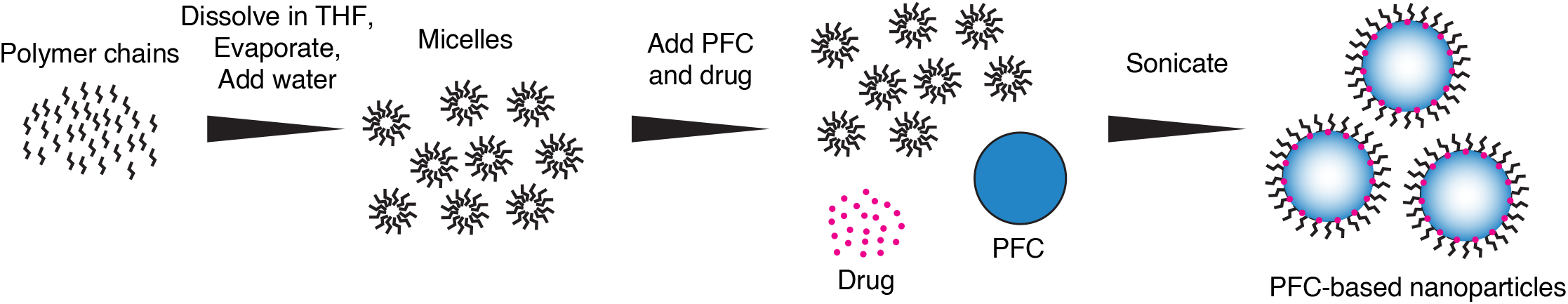
Nanodroplet Production. The conversion of polymeric micelles into PFC-core-based nanodroplets using ultrasound. The three steps are described in detail in the text.

### 5.3 Hydrophobic drug preparation

The propofol used was already in its hydrophobic form, but mycophenolate motefil and ketamine had to be modified from hydrochloride salts to be encapsulated in the nanodroplets. To do so, mycophenolate motefil was first dissolved in deionized water in a centrifuge tube at a concentration of 10 mg/mL. Ketamine was obtained in solution at a concentration of 100 mg/mL. Then, 3N NaOH was added dropwise at an equal molar ratio with the HCl until the drug precipitate forms. The solution was then centrifuged for 5 minutes at 600 RCF and the supernatant discarded. Centrifugation was repeated for a total of three cycle, resuspending in deionized water after each time. After the final centrifuge cycle, the drug was dissolved in methanol then filtered through a 0.2 micron syringe filter. Methanol was then dried under a gentle nitrogen stream, leaving the hydrophobic drug in a powdered form which could be incorporated into nanodroplets.

### 5.4 Nanodroplet characterization

The sizes were measured using a Zetasizer Nano S (Malvern Panalytical, UK), which reports the intensity-weighted size distribution. The size values reported in the Results section describe the mean ± standard deviation of the distribution of the intensity values measured by the device. To quantify the amount of drug encapsulated, a 50 *μ*L solution of nanodroplets is added to 450 *μ*L of methanol to dissolve all components (14). A UV-Vis spectrophotometer (NanoDrop 2000, Thermo Scientific) is used to quantify the concentration by comparing the absorbance to a standard curve at 276 nm for propofol and 305 nm for mycophenolate motefil.

### 5.5 Apparatus

As in a previous study (17), drug release is quantified in standard 1.5 mL microcentrifuge tubes. Each tube with freshly produced nanodroplets is placed into a plastic holder. A focused ultrasonic transducer (H-115, 64 mm diameter, 52 mm focal depth, Sonic Concepts) was positioned 52 mm below the holder so the sample was within the ultrasound focus. Degassed water (AIMS III system with AQUAS-10 Water Conditioner, Onda) mediated coupling between the ultrasound face and the vial. The transducer was operated at 300 kHz and the third harmonic, 900 kHz. Stimuli were generated using a function generator (33520b, Keysight, USA). The signals were amplified using a 55-dB, 300 kHz–30 MHz power amplifier (A150, Electronics & Innovation, USA).

### 5.6 Ultrasound parameters

The ultrasound carrier frequencies for *in vitro* experiments were 300 kHz and 900 kHz. For the assessment of different PFC cores and drug encapsulated, continuous pulses 100 ms in duration were repeated once per second for a total of 60 seconds (14, 17). The pressure levels at the vial location, measured in degassed water, were 0, 0.5, 0.9, 1.3, 1.7, 2.1, and 2.5 MPa. During the nanodroplet manufacturing study, ultrasound was applied in 10 ms pulses 10 times per second for one minute. The pressure fields were measured using a capsule hydrophone (HGL-0200, Onda) calibrated between 250 kHz and 40 MHz and secured to 3-degree-of-freedom programmable translation system (Aims III, Onda).

### 5.7 Drug release characterization

100 *μ*L of hexane was placed on top of 200 *μ*L nanodroplet solutions prior to sonication to act as a sink for released drug (14). After 1 minute, 45 seconds of total incubation time, 50uL of hexane was extracted. The amount of dissolved propofol was quantified using UV-Vis spectrophotometry as described previously. The percent release efficacy is defined as the amount of the drug released into the hexane relative to the amount encapsulated. Each datapoint in Fig. 2 included 3-4 distinct samples.

### 5.8 Nonhuman primate biocompatibility

Biocompatibility was tested in one rhesus macaque (*macaca mulatta*, male, age 10 years, weight 14 kg). Nanodroplets were administered intravenously once per week using vascular access ports implanted in the right saphenous vein (41). Nanodroplets were manufactured as described above and doses quantified by the concentration of propofol. The dose was ramped up gradually: 0.25 mg/kg, 0.5 mg/kg, and 1.0 mg/kg to detect potential dose-dependent effects. We then reduced the dose to 0.5 mg/kg for three additional sessions as we found this level to induce robust behavior effects in a previous study. 2 mL of blood was drawn before the first dose then 1 and 7 days after each dose. Samples were analyzed by IDEXX and normal ranges were provided by the Association of Primate Veterinarians (green areas in Fig. 4 and Fig. S3. To quantify any potential effects, the baseline for each dose was subtracted from the measurements at day 1 and 7. The lowest dose was excluded from this analysis because its efficacy is unknown. To detect effects, we ran two-tailed, one-sample t-tests for each parameter at days 1 and 7 and computed a 95% confidence interval for each result (Table S3).

## 6 ACKNOWLEDGEMENTS

We thank Dr. Natalya Rapoport for comments and input throughout the study, and Sarah Haslam for assistance. We thank Dr. Melanie Graham for implanting the macaque with a vascular access port and Dr. Caroline Garrett for excellent veterinary support. This work was supported by the National Institute of Neurological Disorders and Stroke, grants R00NS100986 and 1RF1NS128569.

## 7 AUTHOR CONTRIBUTIONS

J.K. and M.G.W. designed the studies. M.G.W. synthesized the nanodroplets and carried out core and frequency drug release and stability experiments. A.P. and M.G.W. carried out manufacturing experiments. A.D. and M.G.W. collected the blood samples and analyzed the resulting data. A.B. optimized methods to synthesize ketamine nanoparticles. J.K. and M.G.W. wrote the manuscript and all authors contributed to editing.

## 9 DATA AVAILABILITY

All data are shown in the respective plots. Raw data will be made available on reasonable request.

## SUPPLEMENTARY MATERIAL

**Table S1.** Summary of nanodroplet core and ultrasound effects. The effects of the nanodroplet core, ultrasound frequency (*F*), and ultrasound pressure (*P*). These effects were assessed using a three-way ANOVA that featured the three main effects and all possible interactions. Bold entries are significant (*p <* 0.05).

| Factor | Significance |
| --- | --- |
| Core | < **0.001** |
| *F* | < **0.001** |
| *P* | < **0.001** |
| Core × *F* | < **0.001** |
| *F* × *P* | < **0.001** |
| Core × *P* | **0.030** |
| Core × *F* × *P* | **0.049** |

**Table S2.**
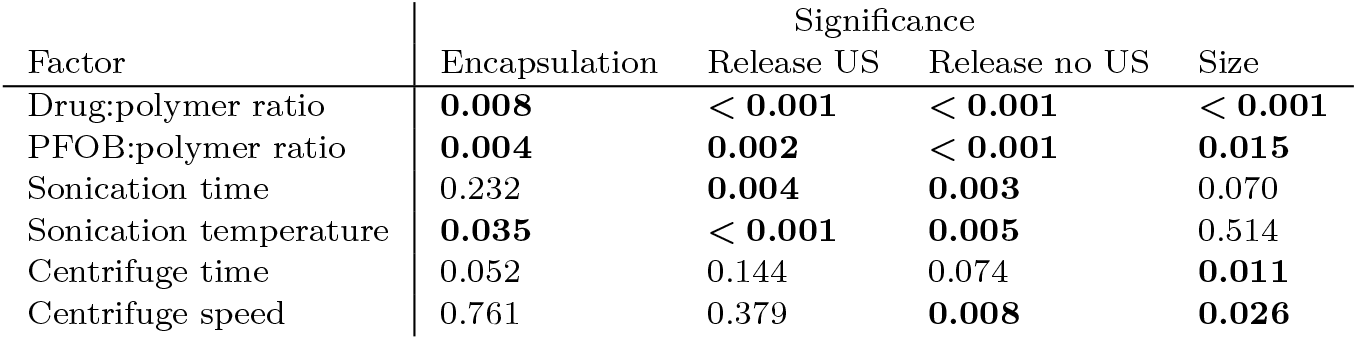
Summary of manufacturing parameters. The effects of nanodroplet manufacturing parameters assessed using a one-way ANOVA for each factor and metric tested. Bold entries are significant (*p <* 0.05).

| Factor | Significance |  |  |  |
| --- | --- | --- | --- | --- |
|  | Encapsulation | Release US | Release no US | Size |
| Drug:polymer ratio | **0.008** | < **0.001** | < **0.001** | < **0.001** |
| PFOB:polymer ratio | **0.004** | **0.002** | < **0.001** | **0.015** |
| Sonication time | 0.232 | **0.004** | **0.003** | 0.070 |
| Sonication temperature | **0.035** | < **0.001** | **0.005** | 0.514 |
| Centrifuge time | 0.052 | 0.144 | 0.074 | **0.011** |
| Centrifuge speed | 0.761 | 0.379 | **0.008** | **0.026** |

**Figure S1.**
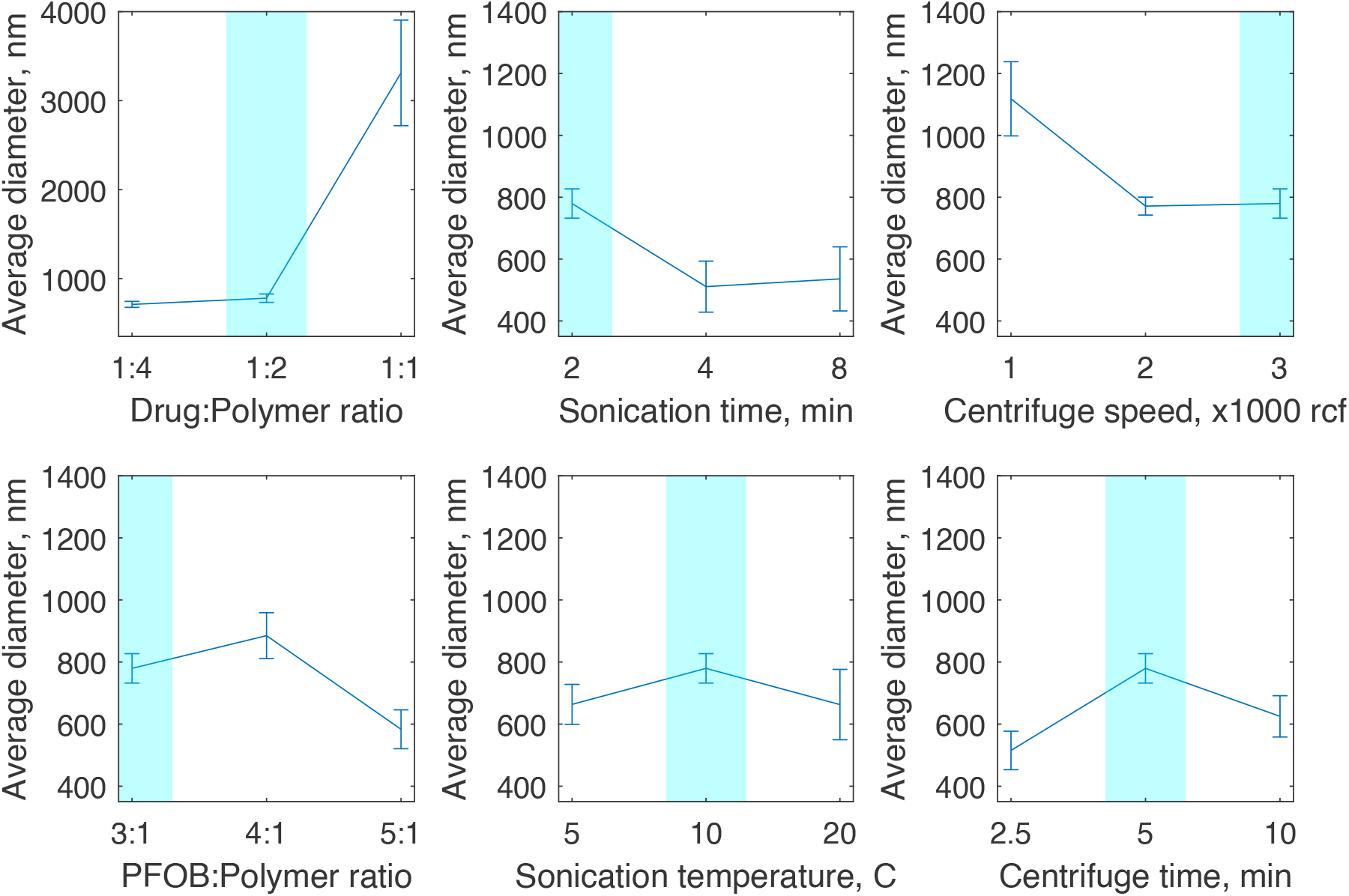
Effects of manufacturing parameters on particle size. Mean±s.e.m. average diameter of nanodroplet samples for each manufacturing method.

**Figure S2.**
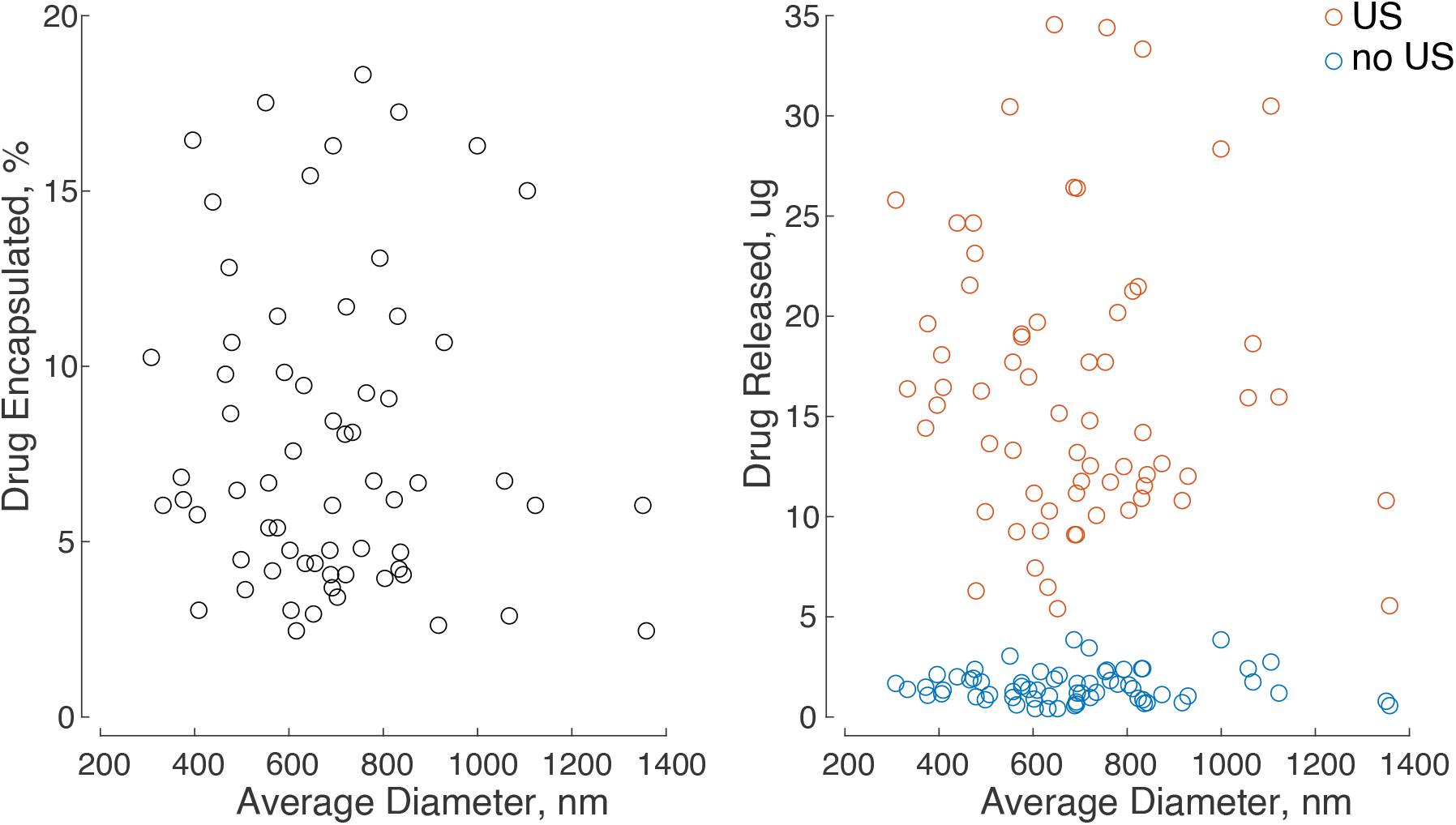
Particle size effects on drug encapsulation and release. Left: Drug encapsulation in nanodroplets as a percentage of drug used for each sample in the manufacturing experiments. There is no significant correlation between average particle size and drug encapsulated. Right: Drug released from each nanodroplet sample with (orange) and without (blue) ultrasound (US). There was no correlation between particle size and drug release either with or without US.

**Figure S3.**
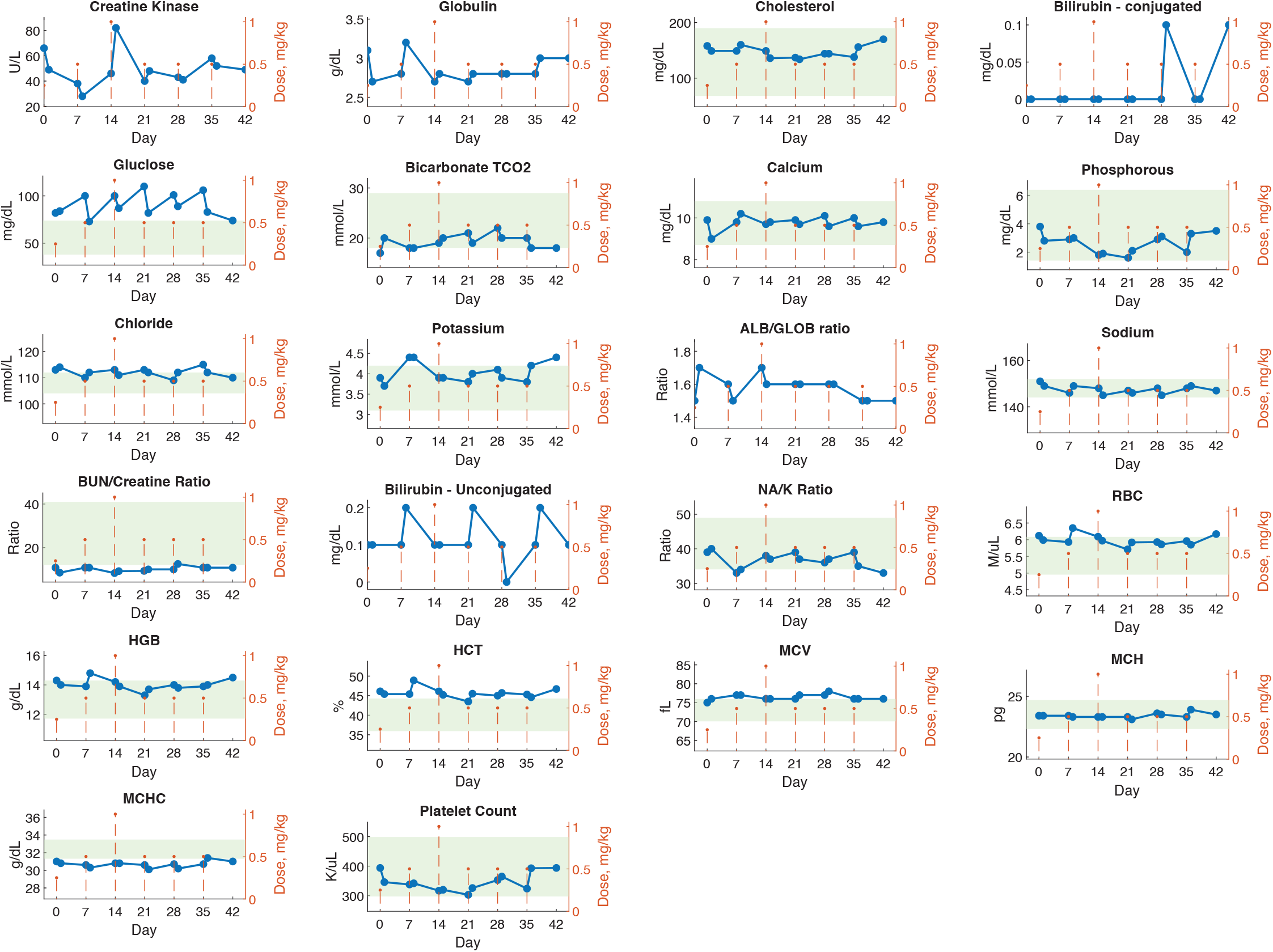
Blood chemistry and hematology from 6-week nanodroplet dosing. Blood chemistry and hematology values for one macaque monkey plotted over 42 days with 6 nanodroplet doses (orange dotted lines). Blood samples were collected one and 7 days after each dose (blue points). Green shaded areas represent normal values established by the Association of Primate Veterinarians.

**Figure S4.**
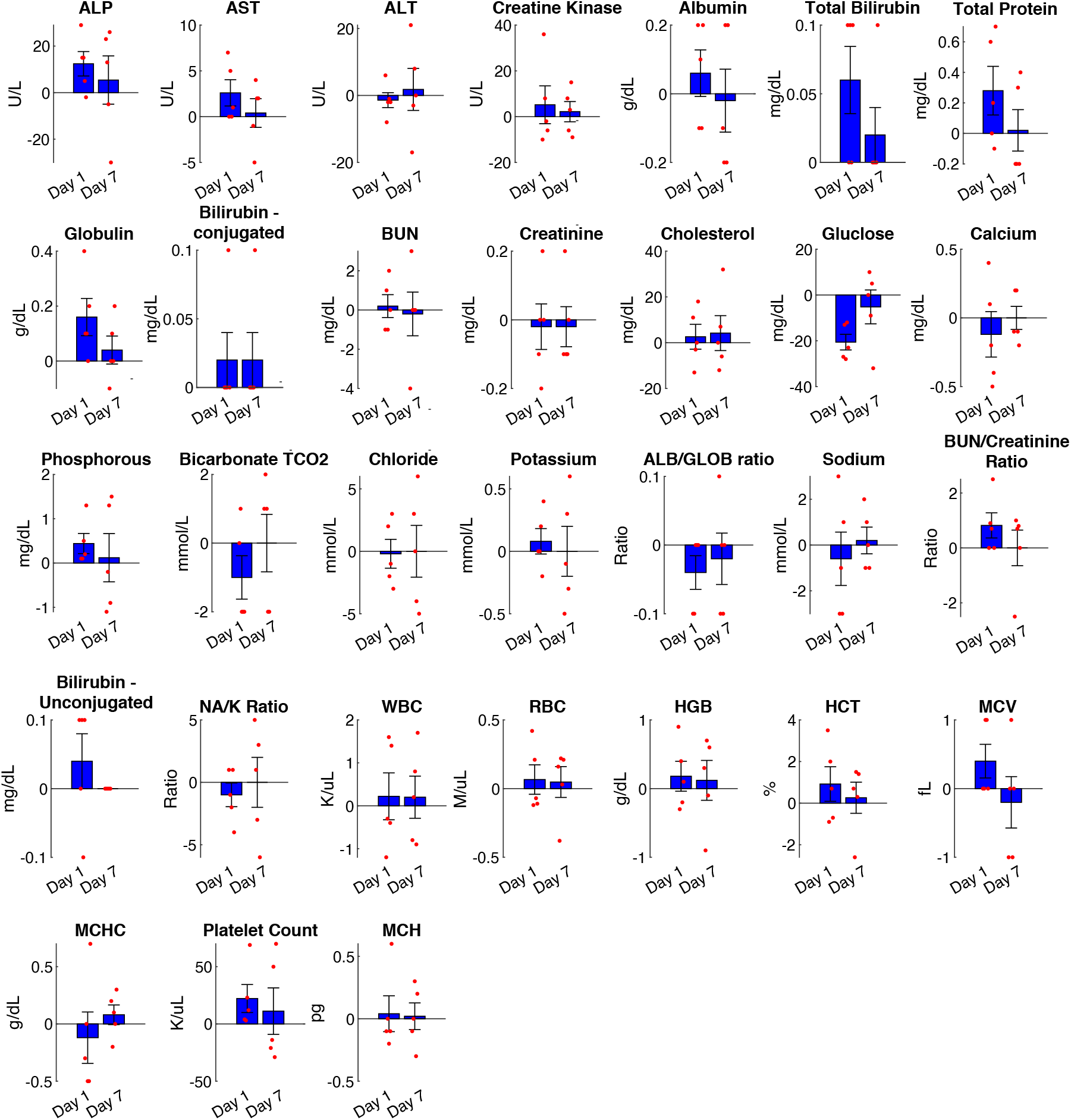
Blood chemistry and hematology values one and seven days after nanodroplet dosing. Change in values, relative to baseline, plotted as individual datapoints (red points) and mean±s.e.m. (blue bars). Results from the five nanodroplet doses at or above 0.5 mg/kg also shown in Fig. 4 and Fig. S3. Only the one-day change in glucose was statistically significant (Table S3; see text for details).

**Table S3.**
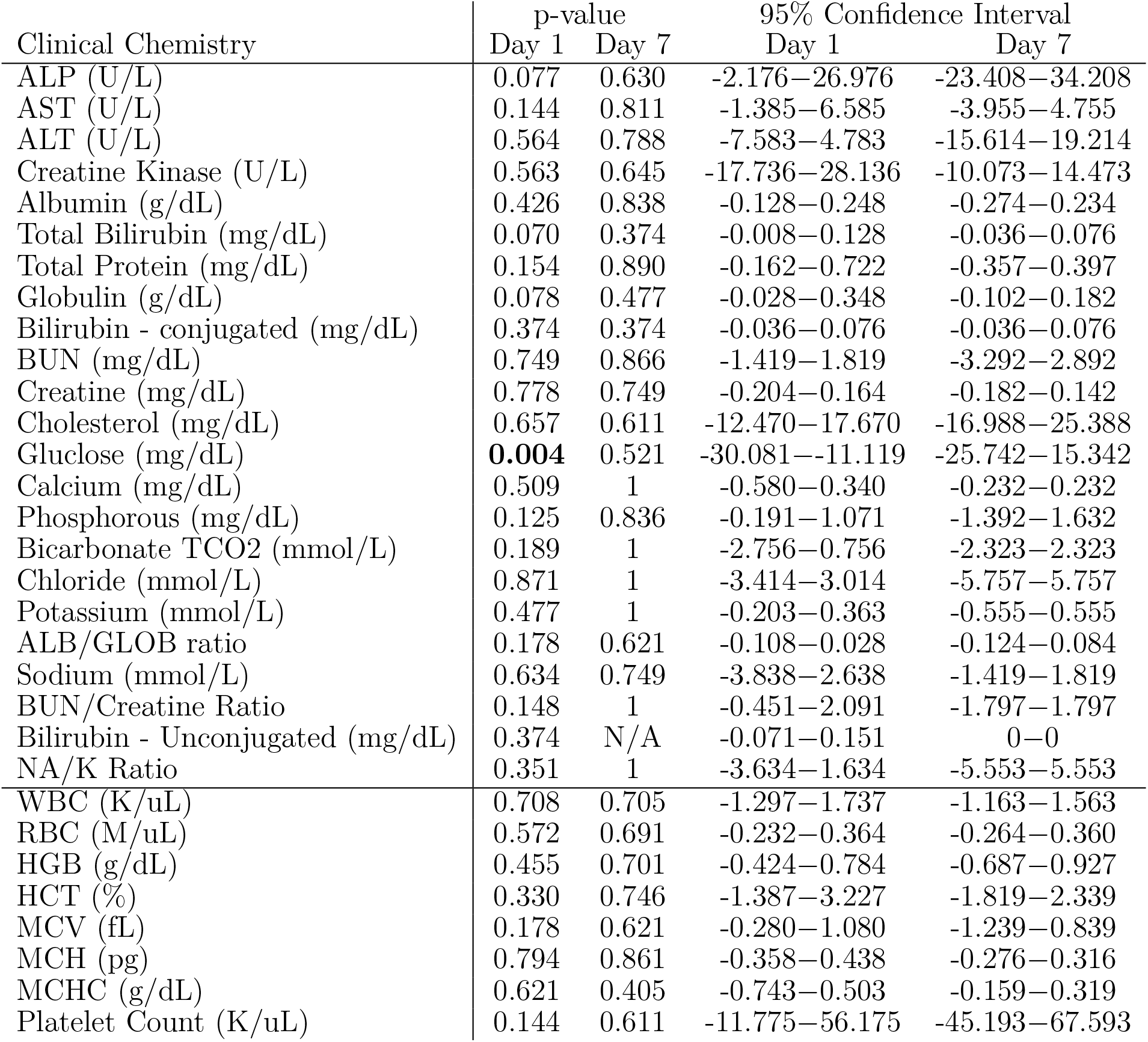
Statistical Analysis of nanoparticle blood compatibility. P-values represent the results of one-sample t-tests of the data shown in Fig. S4. *n* = 5 for each test. Bold entries are significant (*p <* 0.05). Confidence intervals indicate the expected range of the change in each value as a result of nanodroplet administration.

